# Cross-species Functional Alignment Reveals Evolutionary Hierarchy Within the Connectome

**DOI:** 10.1101/692616

**Authors:** Ting Xu, Karl-Heinz Nenning, Ernst Schwartz, Seok-Jun Hong, Joshua T. Vogelstein, Damien A. Fair, Charles E. Schroeder, Daniel S. Margulies, Jonny Smallwood, Michael P. Milham, Georg Langs

**Author notes:** These authors contributed equally to this work (Ting Xu, Karl-Heinz Nenning). These authors contributed equally to this work (Michael P. Milham, Georg Langs).

## Abstract

Evolution provides an important window into how cortical organization shapes function and vice versa. The complex mosaic of changes in brain morphology and functional organization that have shaped the mammalian cortex during evolution, complicates attempts to chart cortical differences across species. It limits our ability to fully appreciate how evolution has shaped our brain, especially in systems associated with unique human cognitive capabilities that lack anatomical homologues in other species. Here, we demonstrate a function-based method for cross-species cortical alignment that leverages recent advances in understanding cortical organization and that enables the quantification of homologous regions across species, even when their location is decoupled from anatomical landmarks. Critically, our method establishes that cross-species similarity in cortical organization decreases with geodesic distance from unimodal systems, and culminates in the most pronounced changes in posterior regions of the default network (angular gyrus, posterior cingulate and middle temporal cortices). Our findings suggest that the establishment of the default network, as the apex of a cognitive hierarchy, as is seen in humans, is a relatively recent evolutionary adaptation. They also highlight functional changes in regions such as the posterior cingulate cortex and angular gyrus as key influences on uniquely human features of cognition.

## Introduction

The human brain differs from other species in both scale and organization^1–5^. One way to understand the functions of this highly complex system is through comparative neuroimaging studies across species^3,6–9^. Evidence of similarities between neural systems in different species suggests a function may be relatively conserved across evolution^10,11^. In contrast, regions showing the greatest changes between humans and other species may highlight neurocognitive systems that are relatively unique to our species^12,13^. Traditionally, cross species comparisons and alignment have depended on the identification of anatomical anchors (e.g. key cortical landmarks or common white matter tracts) and corresponding cortical features (e.g. myelination) in humans versus other species^3,14–17^. This approach has allowed comparing human cortex to non-human primates including macaques, marmosets and chimpanzees, leading to the identification and comparison of putative ‘homologue’ regions across species^14,15,18,19^. This use of anatomical features has revolutionized our understanding of the organization of the mammalian cortex.

An emerging feature of these observations is that brain evolution is not a globally homogeneous phenomenon (e.g., a simple scaling), but a more complex process of reorganization (e.g. mosaic processes)^1,4,20^. Unimodal cortical regions, that play well-defined roles in perception and action, are preserved and remain relatively consistent across species, suggesting basic processes linked to moving and perceiving are relatively conserved across species^10,11^. In contrast, relatively large evolutionary changes in cognitive capabilities may have emerged in processes that are less closely tied to the processing of information in the ‘here and now’^19,21–24^. These include the ability to understand that conspecifics hidden mental states that can influence our behavior (i.e. theory of mind), to solve problems in a creative manner, to use language to communicate intentions, and, to explicitly use imagination to consider different times and places^25–29^. In humans, many of these processes are related to neural processing in regions of the so-called default mode and frontoparietal networks^30,31^. These systems are relatively distinct from unimodal regions, and often lack clear anatomically-defined cross species homologues^12,14^, limiting our ability to provide a precise understanding of their similarities or differences.

A fundamental challenge in understanding the evolution of cortical organization, therefore, is how to extend our understanding of cross-species differences in neural function to regions of association cortex where there are relatively few well-defined physical landmarks – and where we may anticipate the largest cross-species differences. Contemporary accounts suggest that mammalian neural processing is organized along multiple hierarchies that describe how information from distinct neural populations are integrated and segregated across the cortex^12^. One of the more important hierarchies reflects the process through which information from unimodal systems are bound together to form abstract, cross-modal, representational codes assumed to be important for multiple aspects of higher order cognition^32^. Importantly, recent observations based on fMRI data have demonstrated this hierarchy is reflected in the geodesic structure of the cortex such that regions of association cortex occupying locations equidistant from unimodal systems (e.g. visual, auditiory, and sensorimotor cortices)^30^. Positioning a region (e.g. the association cortex) to be equidistant from unimodal cortices could allow its neurons to respond in a relatively unbiased fashion to information from multiple modalities. This in turn would provide a mechanism that allows neural representations to extend across modalities. Studies have shown that at a coarse level of analysis, similar hierarchies exist across different primate species while the evolutionary changes in association cortex have been more pronounced with greatest expansion^3,14,18^. Converging evidence, therefore, highlights the need to understand how neural hierarchies that span unimodal to transmodal cortex have changed across humans and other species of primate in order to gain a more complete picture of how evolution has shaped cortical function, cognition and behavior.

To provide a more comprehensive understanding of how evolution has shaped cortical functions, therefore, our study leverages recent advances in representing functional organization in a high-dimensional common space^30^. We build on prior work that shows it is possible to align across species by comparing neural activity during movie watching^8^. Our study capitalizes on advances in task-free functional connectivity data collected across two species (human and macaques)^33^. Our method, which we refer to as ***joint-embedding***, simultaneously extracts common dimensions of shared functional organization from both humans and macaques. It establishes a common coordinate space for both species in which comparisons of functional organization between species can be made in a quantitative manner. Using this approach, we first examined whether the joint-embedding approach can identify the cross-species landmarks in a manner that is consistent with what is previously known. Having done so we next explored whether our approach can shed light on how changes in the hierarchical organization of unimodal and transmodal regions across species, may contribute to the emergence of uniquely human features of higher order cognition.

## Results

To promote replication and reproducibility all of the methods presented here were obtained from openly available resources. Specifically, the Human Connectome Project (HCP) S500 release (n = 178) and the PRIMatE Data-Exchange (PRIME-DE)^33,34^. From the latter, three cohorts were selected for analysis - two of which were anesthetized samples, from the Oxford University (n=19, 53.3 min per monkey) and UC-Davis University (UCD, n=19, 6.7 min per monkey), and one awake sample from Newcastle University (n=10, 8.3 min per monkey)^35–39^. We randomly divided HCP into two subsets to construct two human and anesthetized macaque comparisons (HCP1-Oxford, HCP2-UCD) and two human and awake macaque comparisons (HCP1-Newcastle, HCP2-Newcastle). Interspecies alignment analyses were replicated in all four comparison samples to ensure their replicability. We focus on the HCP1-Oxford sample in the main results and present the three other comparisons in supplementary material.

### Joint-embedding approach captures the common brain architecture across species

To construct a common functional space for cross-species comparison, we extended the spectral embedding-based approach for mapping connectivity topographies (termed as ‘connectopies’ in short)^30,40–42^. Specifically, rather than computing spectral embeddings for each species individually and subsequently performing alignment, we applied embedding to a joint similarity matrix (Fig 1A, details in Method). The joint similarity matrix was constructed by concatenating the following four submatrices: 1) two within-species similarity matrices (one for each species located along the diagonal), calculated using cosine similarity of thresholded functional connectivity at each vertex in each species, and 2) two off-species similarity matrices (macaque-to-human and its transpose human-to-macaque), calculated by cosine similarity of functional connectivity at each vertex with each of matched homologous landmarks, and treated as off diagonal matrices (Fig 1B, Table S1 and Method)^14,43–45^. The joint-embedding results in a representation of functional connectivity shared between the two species in the form of components. Specifically, we extracted the top 200 matched components for both species, with each dimension represents the common feature along the respective embedding axis. We refer to this common space as the joint-embedding space and each dimension (component) as a ‘gradient’. Of the 200 gradients, the top 15 components met our minimum landmark alignment criteria and were retained for further analysis (see Methods). We validated the matched inter-species common space by examining the similarity of the presumably anatomical landmarks in this space (Fig 1C, Fig S1). The Pearson correlations of the landmarks in joint-embedding space are all in top 5% percentile of the pairwise correlations between human and macaque (ranging from r=0.877 [FEF] to r=0.999 [V1], p<0.001; Fig S1).

**Figure 1.**
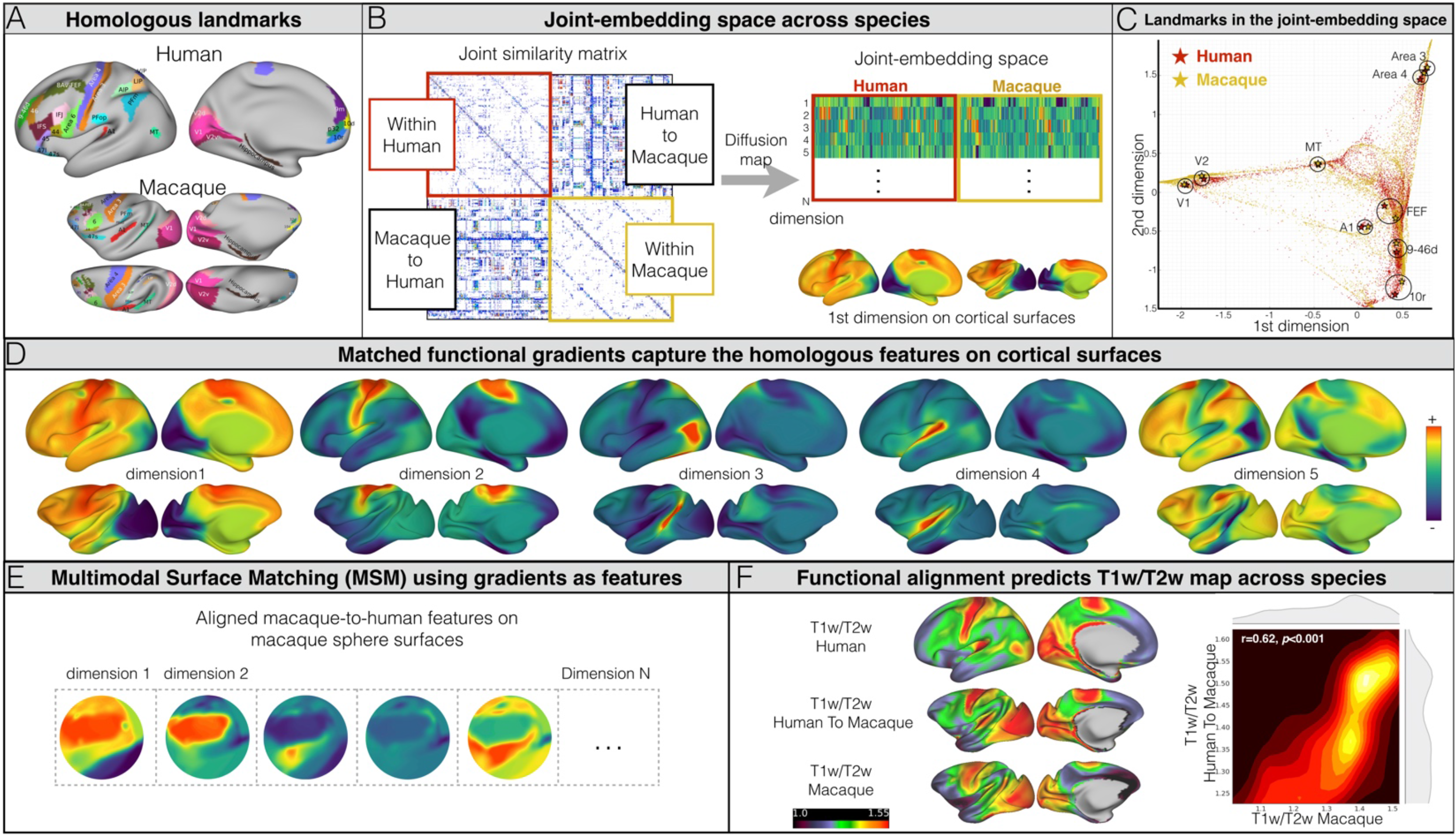
Joint-embedding captures the common brain architecture between human and macaque monkey. A) Cross-species homologous landmarks defined by previous studies (Supplementary Table S1). B) Schematic diagram for constructing the cross-species functional common space. The joint-similarity matrix is concatenated by the vertex-wise within-species similarity matrix (diagonal) and between-species similarity matrix (off-diagonal). Spectral embedding was applied on the joint similarity matrix to extract N number of the matched components to construct the cross-species common space. C) Homologous landmark pairs are close in the joint-embedding space (Supplementary Fig S1-2). D) Matched components (i.e. gradients on cortex) on human and macaque cortical surfaces highlight the homologous areas. E) The gradients were used as features in MSM for cortical surface alignment on sphere surfaces. F) The established alignment can predict the T1w/T2w map based on the other species. As visualized in a 2D density plot, the T1w/T2w in macaque (x-axis) shows significant spatial correlation with the human-to-macaque T1w/T2w prediction (y-axis).

### The joint embedding method highlights the homologous areas and distribution of myelin and so can serve as a method of cross-species alignment

Our first set of analyses are aimed at validating our cross-species joint-embedding. To demonstrate that the gradients can serve as common space for both species, we show that they capture the similarity in location of the well documented cross species landmarks, and that they also describe the known distribution of myelin. After generating the joint-embedding space we projected the results (i.e. gradients) back onto human and macaque cortical surfaces. As shown in Fig 1D, the homologous areas for both species are the apex of the same gradient (e.g. V1 as a negative nadir in gradient 1, MT+ as the positive apex in gradient 3) or occupy similar areas within the spectrum of a gradient (e.g. MT+ in gradient 1, V1 in gradient 3).

Next, we examined whether the joint embedding gradients provides a compact description of the distribution of myelin in both species. To establish the surface deformation between macaque and human cortex, the top 15 gradients were used as functional mesh features in Multimodal Surface Matching (MSM) for the surface registration between human and macaque (Fig 1E; details of the model are in Methods)^40,46^. We validated the alignment generated by joint embedding by testing its ability to generate a myelin sensitive map for each species though predicting the myelin sensitive map (i.e. T1w/T2w) based on the other species following the established alignment^47^. We applied the surface human-to-macaque alignment to the myelin map that calculated from HCP data and compared the aligned T1w/T2w predicated map to the actual macaque T1w/T2w map calculated based on Yerkes-19 template sample (Fig 1F)^48^. The predicted map was similar to the actual T1w/T2w map (r=0.622, p<0.001). We also applied the macaque-to-human alignment to predict the human T1w/T2w using the macaque T1w/T2w map, yielding a similar, although weaker association (r=0.574, p<0.001).

### The cross-species functional homology index (FH Index)

Having established that our cross species embedding adequately captures both the known homologous landmarks, and species-specific distributions of myelin in a reasonable manner, we next considered whether our approach can provide a description of how the cortical surface differs between macaques and humans. To quantify regional similarities in functional organization across species, we developed the Functional Homology Index (FHI, Fig 2A). For each pair of coordinates identified as corresponding between species using MSM, the FHI quantifies the maximum similarity of functional gradient profiles across species within corresponding searchlights (radius = 12 mm along the surface). An advantage of using a searchlight approach is that it mitigates the possibility of excessive topological constraints from MSM, while limiting the identification of matches that are unfeasible. The maximal similarity within the corresponding searchlight evaluated the highest likelihood that the functional gradients at each vertex in human can be represented in macaque (Fig 2B) and vice versa (Fig S4B). These patterns were replicated in all four independent comparison samples (Fig S5).

**Figure 2.**
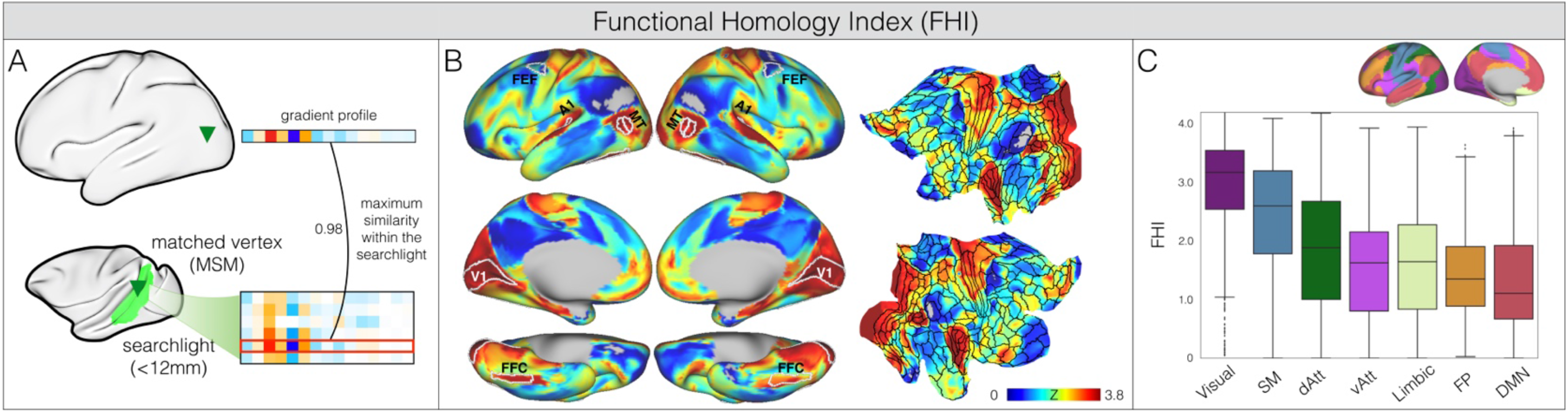
The functional homology index (FHI) reveals the cross-species similarity in network hierarchy. A) FHI is calculated as the local maximum similarity of the functional gradients profile across species within corresponding searchlights (geodesic distance < 12 mm from the MSM matched vertex). B) FHI exhibits high values in sensory cortices (e.g. visual, auditory, somatomotor), and lower in high-order association regions. C) FHI reveals the network hierarchy (Yeo2011 networks).

The FHI revealed the upper bounds of interspecies alignment that can be achieved based on functional organization when going from human-to-macaque, and from macaque-to-human (Fig S4B). Here, the macaque-to-human and human-to-macaque functional homology indices reflect the degree of common functional organization across both species. The cross-species homology index was most prominent in sensory areas including early visual cortices, the motion-selective visual area (MT), auditory area, the fusiform face area (FFC) and the dorsal somatomotor areas. As expected, the prefrontal cortex, which is greatly expanded in human relative to the macaque, exhibited lower a homology index. Importantly, a large set of high-order association regions showed the lowest degree of homology index, in particular in dorsolateral prefrontal cortex (dlPFC), lateral temporal cortex (LTC), parietal gyrus regions (PG), and posterior medial cortex (PMC). This set of regions corresponds well with what is known as the default mode network (DMN) in human. Of particular interest, the FHI is at zero in PG regions, indicating that no functional corresponding regions can be found in macaques that have a functional organization similar to that of PG regions in humans. To quantify the FHI at the level of networks, we averaged macaque-to-human similarity strength in each of seven human networks (Fig 2C)^49^. The sensory networks, in particular the visual network, showed the highest homology index, followed by the attention, limbic, frontoparietal networks with a moderate degree of homology index; the DMN showed the lowest homology index. Next, we examined the associations between spatial distribution of the FHI and the distribution of myelin. We found that the T1w/T2w myelin sensitivity map was significantly associated with the FHI such that greater similarity between species was linked to greater estimated levels of myelin (Fig S4C, r=0.428, p<0.001).

### The cross-species functional homology index characterizes the distributed local sensory hierarchies

Our analysis so far suggests that the functional organization of the cortex has changed the least in regions of unimodal cortex, and that these tend to be regions that show relatively greater levels of myelin. Next we examined whether the FHI also can describe more local (i.e., within system) variations in cross-species functional organization. Here we focus on examples from the visual and somatomotor systems, as their hierarchical organizations are among the best understood^12,50^.

The distribution of the FHI in the visual system can be seen in Fig 3. It can be seen that in the early visual system, the primary visual area, V1, has the highest homology index, followed by the secondary visual area V2, the third visual area V3, and V3A; the lowest functional homologue values were observed in V4. This pattern appears to reflect previously established differences in the visual eccentricity processing pathways^50^. Beyond early visual areas, in the ventral pathway, we found that the mean functional homology index progressively decreased with the complexity of information across ventral stream labeled by Brodmann Atlas areas: 17, 18, 19, 37, 20, 21^51^. The dorsal system exhibited a more complex pattern. V6 and V6A were found to have higher FHI than V7 and IPS, though these regions did not differ substantially from one another (Fig S6A). Notably, consistent with the tethering hypothesis, which suggests that as V1 and MT were “molecular anchors” for evolutionary processes, the FHI for MT+ was comparable to V1, with lower scores being noted in surrounding areas (Fig 3C). In somatomotor areas, the FHI varied along the dorsal-ventral axis of somatotopic mapping^52^. The lower limb (i.e. foot) area has the greatest homology index, followed by trunk (i.e., body), and upper limb (i.e., hand); The ventral areas, eye and face/tongue areas have a lower homology index (Fig 3E).

**Figure 3.**
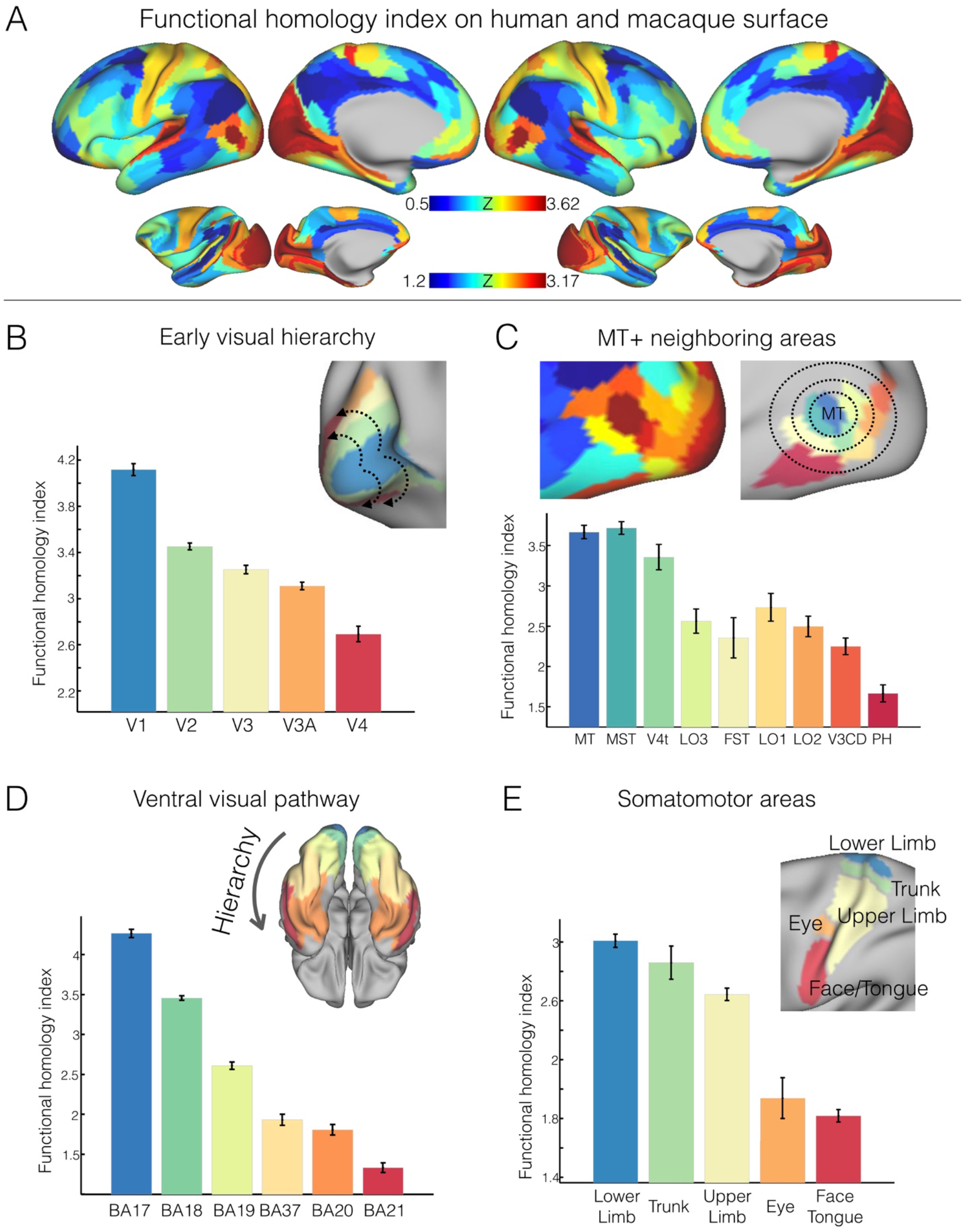
Cross-species functional homology index characterizes distributed local sensory hierarchies. A) Parcel-wise FHI maps in human and macaque monkey based on the recent parcellations (human: Glasser et al., 2016; macaque: Markov et al., 2011). B) FHI reflects the hierarchy in the early visual processing system. Five areas defined by the Glasser parcellation are rendered on the surface and comprise the early visual areas ordered by the hierarchical streams of eccentricity mapping on x-axis (V1, V2, V3, V3A, V4). C) FHI exhibits the highest scores in MT and MST with lower scores in MT+ neighboring areas (x-axis: ordered by the geodesic distance from MT area). D) FHI decreases along the ventral visual processing hierarchy (BA17, BA18, BA19, BA37, BA20, BA 21). E) FHI map varies along the dorsal-ventral axis of somatotopic mapping. The FHI was averaged within labeled areas and visualized with 95% confidence intervals. *Note:* FHI in dorsal visual pathway and primary auditory areas is depicted in Supplementary Fig S6.

### The FHI reveals the modular specialization of the subsystems in attention and frontoparietal networks

Our examination of association areas began with the frontoparietal and attention networks. In humans the frontoparietal and attention networks have been suggested to be dissociable into pairs of networks that are dissociable with respect to their functional proximity to unimodal versus transmodal systems (i.e. the default mode network) ^53,54^. In our analysis, the FHI readily distinguished between the frontoparietal network-A (FN-A) and frontoparietal network-B (FN-B). Specifically, we found that FN-A which has stronger connections to the default network and exhibited lower FHI scores (Fig 4A); while, FN-B, which is more connected to the dorsal attention network in humans^53,54^, exhibited higher scores. Similarly, a lower FHI was observed in dorsal attention network-A (dATN-A) than dATN-B, which is more directly connected to retinotopic visual regions^54^. Finally, a similar pattern was observed in the two subnetworks in ventral attention network. These patterns were replicated in all four independent comparison samples. Within both the fronto-parietal and attention systems, therefore, we found a consistent pattern that the FHI was lower in the sub networks that are functionally less closely linked to unimodal systems^55^.

**Figure 4.**
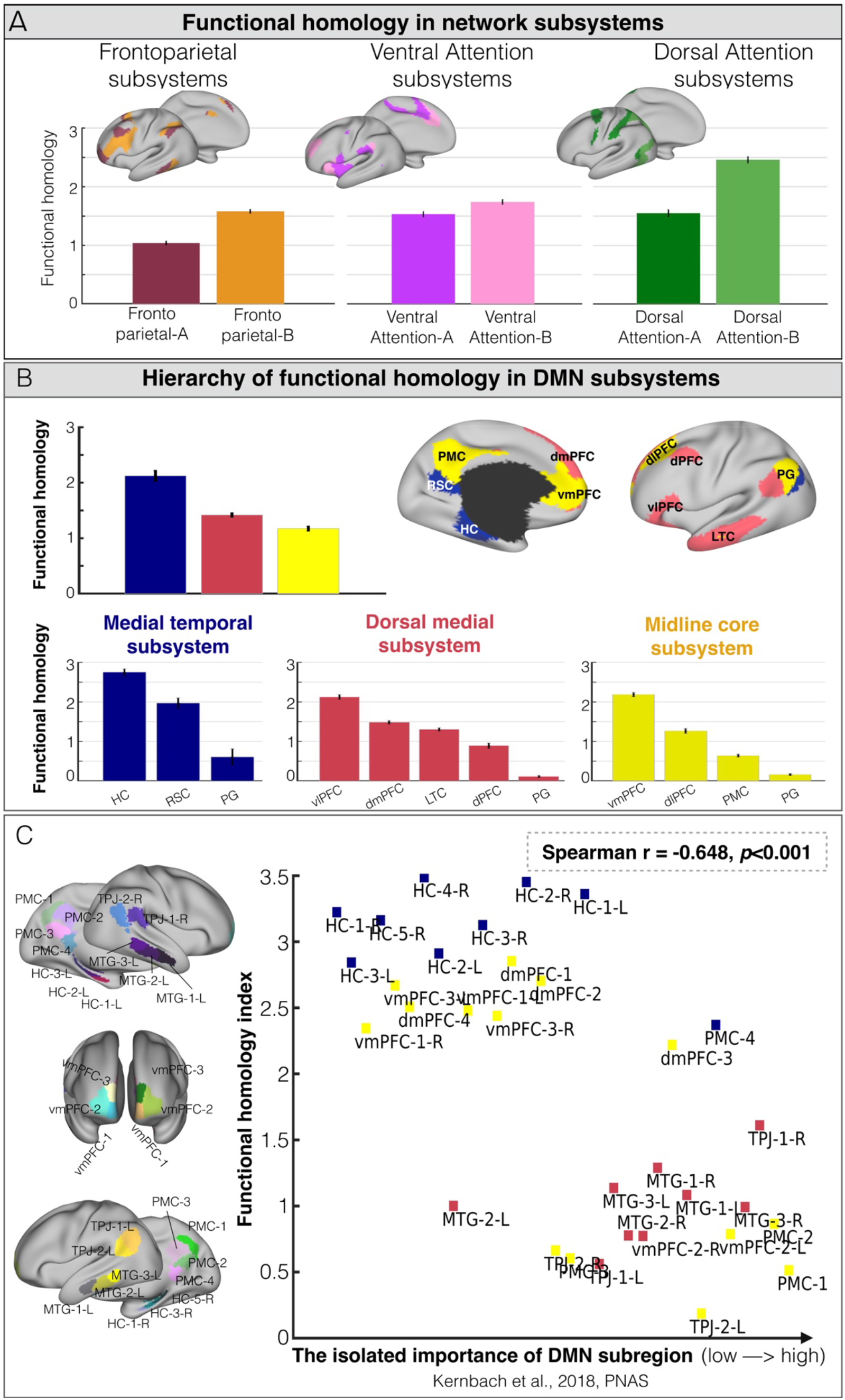
The functional homology index reveals the hierarchy of subsystems in the attention, frontoparietal and default mode networks. A) The FHI shows lower scores in subsystem-A and higher scores in subsystem-B for dorsal attention, ventral attention and frontoparietal networks. B) FHI differs among subsystems of the default mode network; the core DMN shows the lowest score, dorsal medial system an intermediate score, and medial temporal the highest score. Within each of the subsystems, the PG, PMC and LTC have the lowest FHI scores. C) FHI is associated with the level of ‘importance’ across subregions within the DMN (Spearman r=-0.648, p<0.001). The importance of the DMN subregions is established by a recent study based on the UK Biobank data.

### The evolutionary hierarchy of subregions in default mode networks

Next, we examined the cross-species similarity of the DMN, which in humans is located at the apex of the principle cognitive hierarchy^30^. Similar to the fronto-parietal network, the DMN exhibited overall differences in the FHI among its subsystems^31,54,56^, with the medial temporal having the highest score, the core the lowest, and dorsal medial system intermediate (Fig 4B). Across the systems, the angular gyrus (PG), posterior medial cortex (PMC) and the lateral temporal cortex (LTC) had the lowest FHI scores (Fig 4B). To further catalog differences in functional homology across DMN components, we tested for associations with the level of ‘importance’ within the DMN, as established by a recent study that applied a ‘virtual lesions’ procedure to data from the UK Biobank to quantify subregion importance^57^. In particular, this analysis identified regions including the posterior cingulate cortex, the angular gyrus and ventrolateral prefrontal cortex as playing an important role in mirroring distributed patterns of neural function from outside the cortex within the DMN. We found that mean FHI was negatively correlated with importance rank (Fig 4C, Spearman r=-0.648, p<0.001), suggested that the most important DMN subregions in the human had the least functional homology between species. Specifically, dorsal medial system components, which were previously identified as having relatively low importance, were found to have high FHI, while medial temporal system components, which were identified as being of high importance, had low FHI. Consistent with prior work suggesting divisions within the core subsystem with respect to high versus low importance, we found that vMPFC and DLPFC exhibited a higher FHI, while temporal parietal junction (TPJ) and PMC had a lower FHI^58^. Together this analysis suggests that regions where the FHI tends to be relatively lower, correspond to locations which in humans tend to be regions that are most important for reflecting global patterns of functional connectivity within the DMN^57^.

While a number of recent studies have identified a “default-like” transmodal network in nonhuman populations (e.g., macaque, marmoset, rodents), the extent to which this putative network functions in a manner akin to the human DMN remains an open question^59–64^. To gain further insight into functional homologies, we examined the cross-species similarity of DMN subregion functional organization profiles with one another; this was accomplished by calculating the cosine similarity of gradient profiles in the common joint-embedding space. Fig 5A-B illustrates cross-species similarities among DMN subregions that exceed our sparsity threshold (i.e., top 10% of pairwise human-macaque similarity over the entire cortex). Four DMN subregions in macaque (i.e., hippocampus [HC], vmPFC, dmPFC and vlPFC) were found to have functional organizations similar to those in humans. For each of the macaque DMN subregions, we identified its functional similarity with each vertex on the human cortex based on degree of correspondence observed in their functional organization (i.e. gradient profile) (Fig 5C). The macaque-to-human similarity maps seeded in hippocampus, vmPFC, dmPFC and vlPFC exhibited highly similar spatial patterns as human DMN. Importantly, the FHI indicated low cross-species similarity in the macaque PMC (PCC), angular gyrus (i.e. PG), and retrosplenial cortex (RSC).

**Figure 5.**
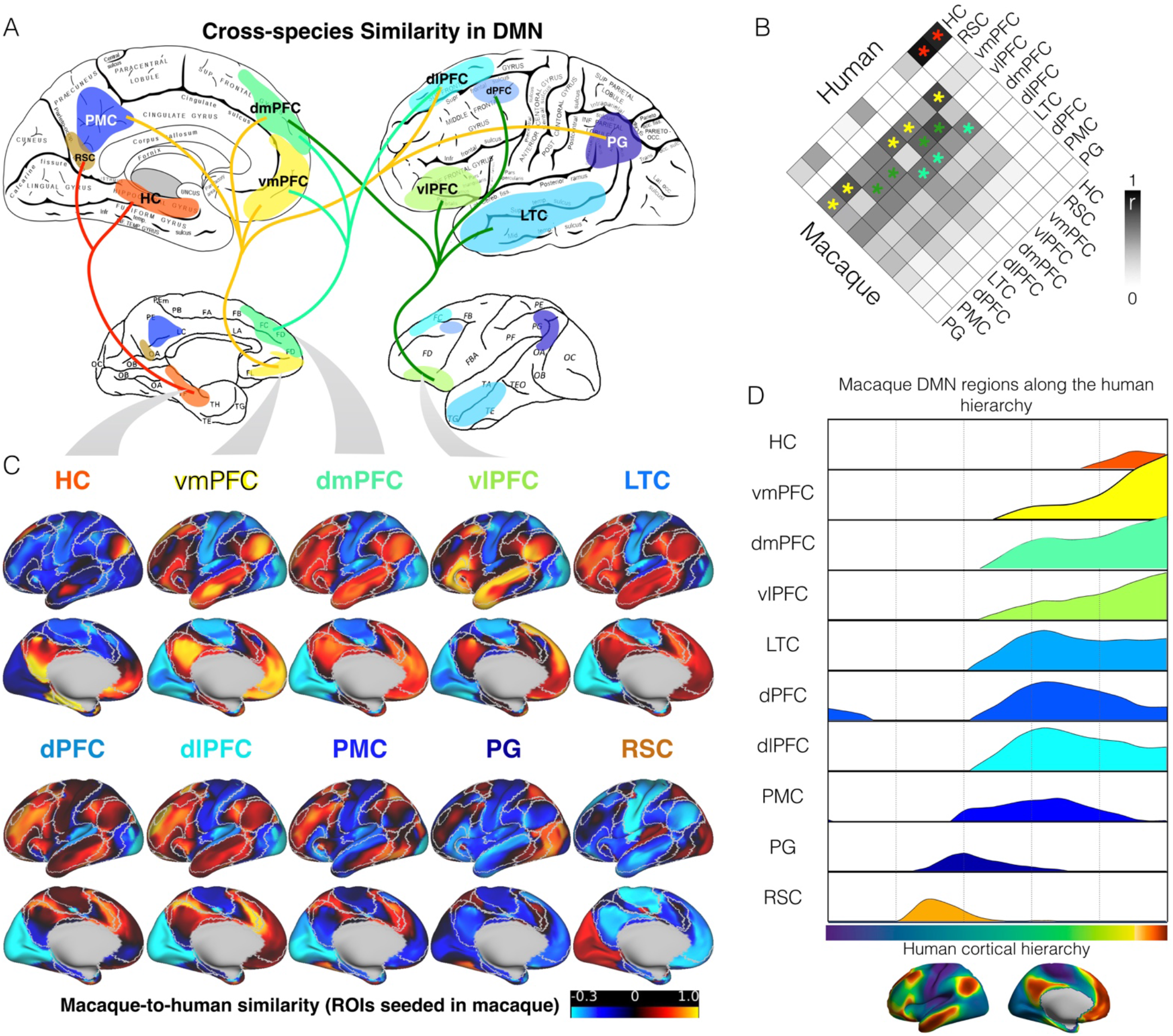
The cross-species similarity between humans and macaque monkeys. A) The human-macaque similarity of functional gradient profiles among DMN candidate subregions. The links in the diagram illustrate the similarity among DMN subregions that exceed the sparsity threshold (top 10% of pairwise human-macaque similarity across the entire cortex). B) The pairwise similarity matrix (cosine similarity) of DMN candidate subregions between humans and macaques. C) The cross-species similarity maps (cosine similarity) for each DMN candidate region seeded in macaque. The macaque-to-human similarity maps of HC, vmPFC, dmPFC and dlPFC regions show highly similar spatial patterns as human DMN (white border based on Yeo2011 networks). D) The macaque-to-human similarity maps are represented along the human principle connectivity gradient obtained based on the human HCP sample. Each line of the macaque DMN seed is smoothed by the locally weighted scatterplot smoothing (LOWESS) kernel. The positive distribution is visualized to explicitly demonstrate the extent to which DMN candidate regions in the macaque reached the human hierarchy apex.

Finally, given recent human studies suggesting the DMN is an apex transmodal network, situated at the furthest end of the macroscale sequence^30^, we examined the extent to which the DMN candidate regions in macaque may have evolved along the cortical processing hierarchy. To accomplish this, we used the human principal connectivity gradient as the reference hierarchy and computed the distribution of macaque-to-human similarity for each of macaque DMN seeds. As shown in Fig 5D, hippocampus, vmPFC, dmPFC and vlPFC in macaque reached the human hierarchy apex, followed by LTC, dPFC and dlPFC. PG, PMC and RSC.

### Cross-species homology maps to cognitive functions

Together, our analysis highlights a pattern of increasing cross species differences in functional organization in regions that are thought to be the most transmodal in humans. Functional imaging studies in humans suggest that these regions are linked to highly abstract, mnemonic states such as semantic and episodic memory, self- relevance, mentalizing or theory of mind, and stimulus independent thought (Margulies et al., 2016). To quantify aspects of human cognition that are often associated with activity within regions of cross species difference, we compared the spatial distribution of the macaque-to-human FHI to those provided by a large-scale meta-analysis of task fMRI experiments^65^. We first grouped the FHI into 10-percentile bins. We then generated probability maps for brain activation under each component and used these to compute the probability strength that each component was associated with activity in each of the 10 bins. Probability strength for each cognitive component was normalized and averaged within each bin (see Methods). Fig 6 shows these data in the form of a heat-map, with the components ordered in rows based on the possibility strengths^30,66^. The higher FHI regions were associated with sensorimotor components (e.g., “visual”, “auditory”, “hand”, and “face”) whereas the lower FHI regions were involved in high-order cognitive functions (e.g. “interoception”, “emotion”, “language”, “reward” and “dorsal attention”). The activation for “working memory”, “inhibition” and “internal mentation” were more likely to overlap with the extremes of the low FHI regions. These findings confirm that the regions in which human neural organization is most different from recent common ancestors are related to a combination of executive functions and introspective processes, both of which reflect aspects of cognition that are assumed to be reasonably unique to our species.

**Figure 6.**
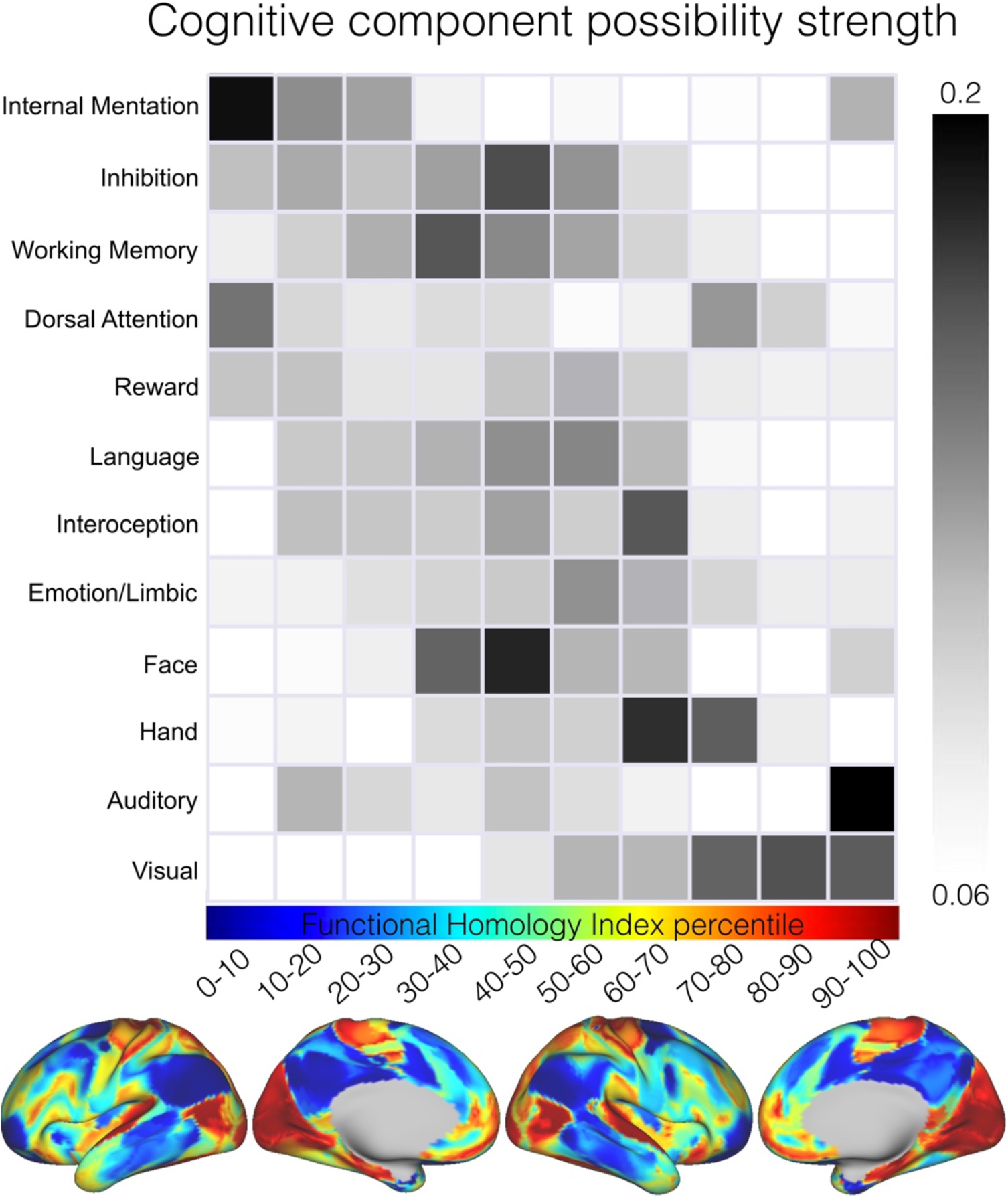
Cross-species functional homology maps to the cognitive functions. Relationship between the FHI map and twelve cognitive components based on the BrainMap meta-analysis database (Materials and Methods). In rows, the percentiles of the FHI map are ordered from low to high. In columns, the cognitive components are ordered based on the normalized activation possibility strength weighted by the log scale of percentile. The higher FHI regions were associated with sensorimotor components whereas the lower FHI regions were involved in high-order cognitive functions.

### The evolutionary surface area expansion and deformation reveals the network hierarchy

So far our analysis has established that regions of maximal cross species difference across humans and macaques, can be understood along a spectrum, from reasonably high levels of similarity in unimodal regions, to relatively low levels of similarity in regions of transmodal association cortex. This pattern was reflected as a functional shift in meta analytic data towards more abstract functions such as working memory or internal mentation that may be considered to be relatively unique to humans^30^. Theories such as the tethering hypothesis attempt to account for how association cortex gains the ability to support abstract functions, by assuming that this emerges from later development in the evolutionary process^12^. To understand whether our index of cross species similarity captures this hypothesized aspect of evolutionary change, our final analysis examined the correspondence between regions that are assumed to have expanded through evolution with the distribution of the FHI. We visualized the relative changes in cortical surface area on the human cortex in Fig 7, in which relatively large. Sensory cortices expanded the least whereas the DMN and frontoparietal network expanded more than 20 times from macaque to human (Fig 7B-C). In particular, the frontal cortex, TPJ, lateral temporal cortex (LTC), and the medial parietal cortex expanded the most from macaque to human. The spatial map of surface area expansion is significantly correlated with the functional homology map (r=-.483, p<0.001) indicating that regions that have expanded the most in human are also those with the least cross species similarity as defined by the FHI. To demonstrate the evolutionary expansion direction more explicitly, we visualized the macaque-to-human deformation vectors on the human inflated surface (Fig 7A, center). In this figure the arrows describe the direction of change, and the color represents the degree of deformation. From macaque to human, the substantial expansion of the frontal cortex appears to push the parietal central regions in a posterior direction. The expansion of TPJ and LTC forced the temporal cortex move posteriorly and occipital visual cortex is then squeezed into the medial side from macaque to human.

**Figure 7.**
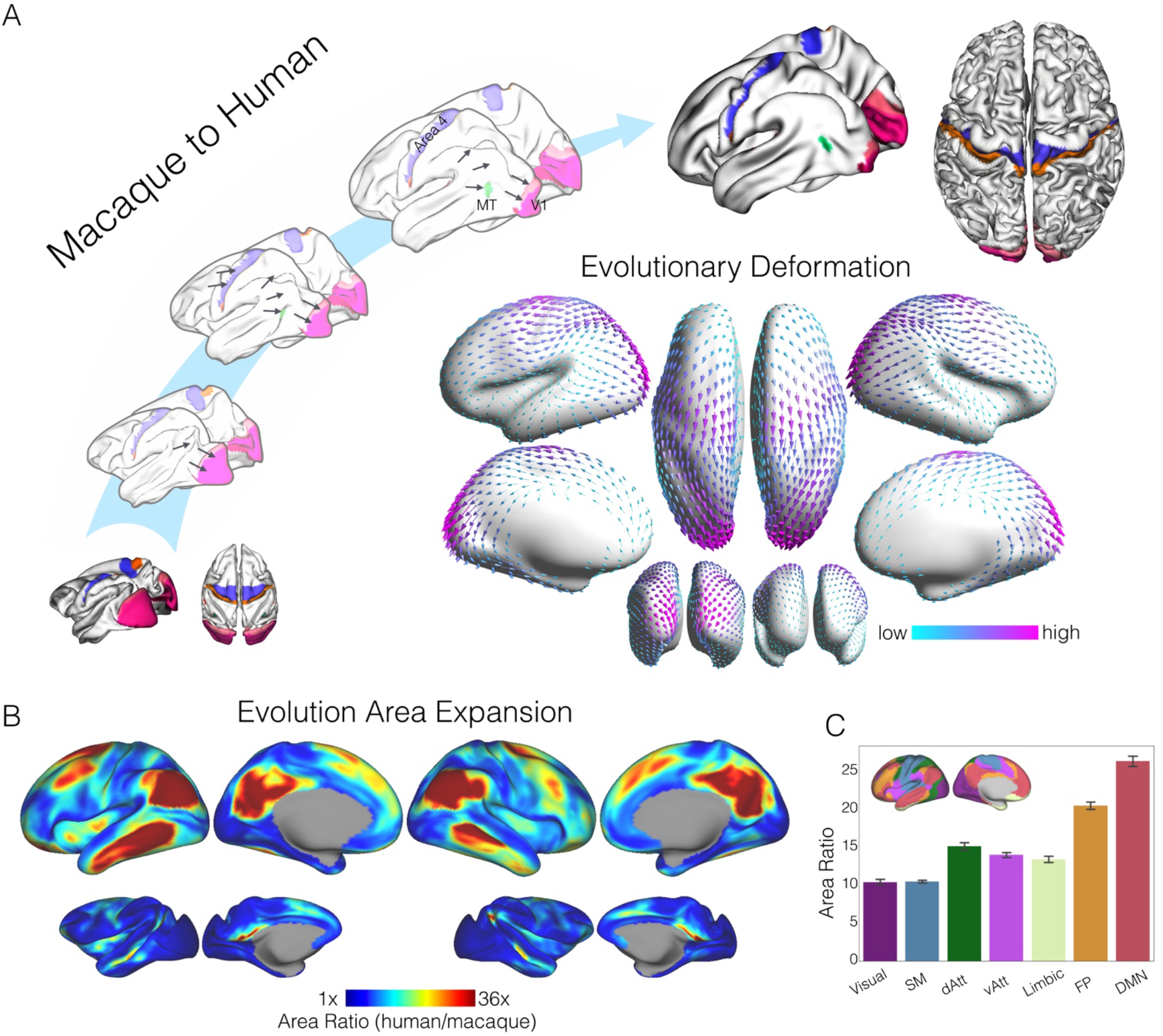
Evolutionary surface area expansion and deformation reveals the network hierarchy. A) Schematic diagram illustrating the evolutionary expansion from the macaque monkey to the human. Homologous anchors in somatomotor (area 3, area 4), primary visual areas (V1, V2) and MT areas were labeled on individual macaque and human surfaces, as well as the intermediate surfaces from macaque to human. Evolutionary expansion direction (i.e. macaque-to-human deformation vectors) is visualized in arrows on the human inflated surface (center). B) Surface areal expansion maps were calculated as the human area divided by macaque area at each of corresponding vertex on human and macaque surfaces. C) Sensory cortices expanded the least (10 times), whereas the high-order association cortex expanded the most from the macaque to the human.

### Addressing a potential confound

In considering our findings regarding the distribution of FHI in sensorimotor areas, one question that may arise is whether the findings may be driven, at least in part, by the specific landmarks included for anatomical alignment and formation of the connectivity matrix upon which joint embedding is based. In particular, whether decreases in FHI as one moves away from the landmark can be artifactual in nature. To address this concern, we repeated our analyses including broader regions in the landmark set, that now include neighboring areas as landmarks in each of FHI hierarchical tests. We found that the hierarchical distributions of FHI reported above remained largely intact (Fig S6). The one exception was for MT+, where differences relative to surround areas were reduced to trend level. It is important, therefore, to treat findings related to MT+ as preliminary. Additionally, for the transmodal default network, where landmarks are more scarce, we added all default candidate areas as a set of landmarks (dmPFC, dlPFC, LTC, PMC) within the network to evaluate potential dependencies for findings regarding this network^67^. As demonstrated in the Fig S6, all DMN relationships reported above were relatively unchanged.

## Discussion

The relative absence of well-recognized landmarks in association cortex between humans and nonhuman primates has important conceptual consequences for understanding how evolution has shaped cortical processing and hence human cognition. In this paper we demonstrate that joint-embedding in a common functional space provides a novel solution to this methodological challenge and reveals important clues as to how and why human cognition may differ from our close evolutionary ancestors.

Joint embedding provides a novel solution to the longstanding challenge of how to align cortical surfaces from different species in order to understand how they have been shaped by evolution. To quantify the likelihood that anatomically homologous areas share a common functional organization across species, we developed the functional homology index (FHI). At a global level, the topography of FHI for humans and macaques appeared to reflect previously identified large-scale cortical hierarchies (meta-analysis based cognitive activation, myelin map and the evolutionary area expansion) with greater similarity in unimodal regions and lower similarity in systems linked to attention and more complex aspects of higher order cognition^12,13,68,69^. A more fine-grained analysis revealed important distinctions were present within canonical circuits in visual and sensory-motor territories^12^. In these areas of cortex, we observed that the landmarks tend to fall in regions that were a local maximum for cross species similarity, and that cross-species similarity declines in adjacent regions. This underscores the value of the landmark approach for identifying how systems directly concerned with input and output systems vary across species, while simultaneously highlighting the value of joint embedding as a means to describe similarity in other regions of cortex^14,16^. We next consider the significance of our findings for several key theoretical issues in the evolution of human cortical organization.

First, using the FHI as an index of evolutionary conservation, the present study provided clear support for mosaicism in the evolution of the human cortex^1,4,20^. Rather than simply differing across brain “modules” or networks, the FHI varied both between and within modules, notably in both cases indexing a hierarchical relationship. Locally within unimodal visual and sensorimotor cortical systems the FHI decreased away from points of well documented correspondence across species^12,70^. Globally the FHI increased from the networks anchored in unimodal cortex and decreased in networks that are important higher order functions^30,71^. Within the DMN those regions with low FHI are the same regions that in humans contain the most information about global brain dynamics. Together these results are consistent with a complex view of cross species differentiation which impacts on cortical modules at both a local and global scale^17,66,68,71,72^.

Second, the distribution of cross-species similarities is consistent with the tethering hypothesis. We found that both the early visual cortex and the ventral visual pathway anchored from V1, and the hierarchy organization seen in MT+ and neighboring areas, were reflected as decreases in the FHI. The tethering hypothesis suggests new functional capabilities have arisen through the gradual duplication, budding, and subdivision of brain areas^12^. Specifically, through a process of cortical expansion, large parts of the cortical mantle are postulated to have progressively become untethered from direct roles in input and output system as they become increasingly distant from the constraints of molecular gradients that surrounded key “anchor” regions (e.g., V1 and MT)^3,12,73^. We found that the FHI reflects this hierarchy at both the local and global scale. For example, in canonical circuits in sensory and motor cortex, the observed radial pattern in the FHI supports the view that evolution changes are increasingly important outside of primary cortex. We also found the FHI was lower in regions in which cortical expansion across species is thought to be largest^3,16^. Although we found evidence in support of the tethering hypothesis in multiple aspects of our analysis, it is important to note that confirmatory analyses could not remove the possibility that the pattern linked to MT may be introduced by inclusion of MT in the landmark set and so this specific finding should be treated as tentative.

Third, our study provides important insight into the evolution of the commonly described DMN as is seen in humans. We found different nodes within the DMN vary in their cross-species similarity. In particular, while some DMN regions are functionally similar in both species, two core regions of the DMN exhibited low similarity between human and macaques: the PMC (PCC-PCU) and PG (parietal gyrus and angular gyrus). In contrast, the vmPFC shows a more similar organizational profile across species^62,74^. Together these results suggest that in functional terms, some aspects of what is called the default mode in humans are present in macaques, while others, particularly the posterior regions, have a much more unique functional profile in humans^31,57^. One implication of the distribution of the FHI across the DMN is that while the foundations of this system may be present in many species, the canonical pattern seen in humans, likely emerged relatively later in evolution. As a consequence, the canonical DMN seen in humans, would be present in only a subset of our recent ancestors. It is also possible that these, as well as higher nonhuman primate species (e.g., champanzee) would exhibit transitional or intermediate DMN variants.

There are a number of issues that should be considered when interpreting our results. Our cross species joint-embedding is part of an emerging trend toward the use of common high dimensional spaces as a tool to understand the evolutionary changes across species. Recent efforts have developed strategies for alignment based on white matter tracts, or myelin (T1w/T2w) maps^15,16^, as well as task-based activations during movie viewing^8^. We anticipate that the joint-embedding approach used in this paper should be readily extensible to diffusion imaging data and encourage future work in this direction^75,76^. Regardless of which data modality is used, the use of a high-dimensional common space may help characterize functional similarities and differences across species, particularly in areas where clear anatomical landmarks are difficult or impossible to ascertain. In principle, this approach could be extended to multiple species (e.g., chimpanzee, baboon, marmoset etc.) to allow alignment across multiple non-human species including human primates, as well as others mammals, for example rodents. It is also important to note that our analyses were carried out at the group-level. This decision was motivated by our desire to maximize signal to noise ratio, though proper implementation of this method in individual animals will require the additional optimization of methods (e.g., sufficient data collection for individual dataset, generation of individual-specific landmark masks)^39,77,78^. In addition, while the present work addressed issues regarding the reproducibility of findings by replicating analyses in independent samples, our understanding of their functional significance is based on a meta-analysis and so the precise functional meaning of the observed differences remains largely a matter of conjecture^54,79^. In order to fully appreciate the significance of these cross species differences in brain organization for human cognition, it will be necessary to understand how these patterns change across contexts, and, if possible, during active tasks states^80,81^. Understanding how functional patterns change across a common embedded space during periods of task engagement, could provide invaluable insight into how evolution has shaped many important aspects of human cognition^24^.

## Acknowledgments

This work was supported by gifts from Joseph P. Healey, Phyllis Green, and Randolph Cowen to the Child Mind Institute and the NIH (BRAIN Initiative R01-MH111439 to C.E.S and M.P.M.; R24 MH11480602 to M.P.M.; NIH NIBIB NAC P41EB015902, Austrian Science Fund FWF (I2714-B31), and the EU (H2020 765148 TRABIT) to G.L.; ERC Consolidator award WANDERINGMINDS 646927 to J.S.; CNRS PICS Grant 288256 to D.S.M.; R01 MH1112439, and P50 MH109429 to C.E.S.; Oesterreichische Nationalbank to K-H.N.; We would also like to thank the investigative teams from Oxford (J. Sallet, R.B. Mars, M.F.S. Rushworth), Newcastle (J. Nacef, C.I. Petkov, F. Balezeau, T.D. Griffiths, C. Poirier, A. Thiele, M. Ortiz, M. Schmid, D. Hunter) and UC-Davis (M. Baxter, P. Croxson, J. Morrison), as well as the funding agencies that make their work possible (Oxford: Wellcome Trust, Royal Society, Medical Research Council, UK Biotechnology Biological Sciences Research Council; Newcaste: National Center for 3Rs, NIH, Wellcome Trust, UK Biotechnology Biological Sciences Research Council; UC-Davis: NIA).

---

**Figure S1.**
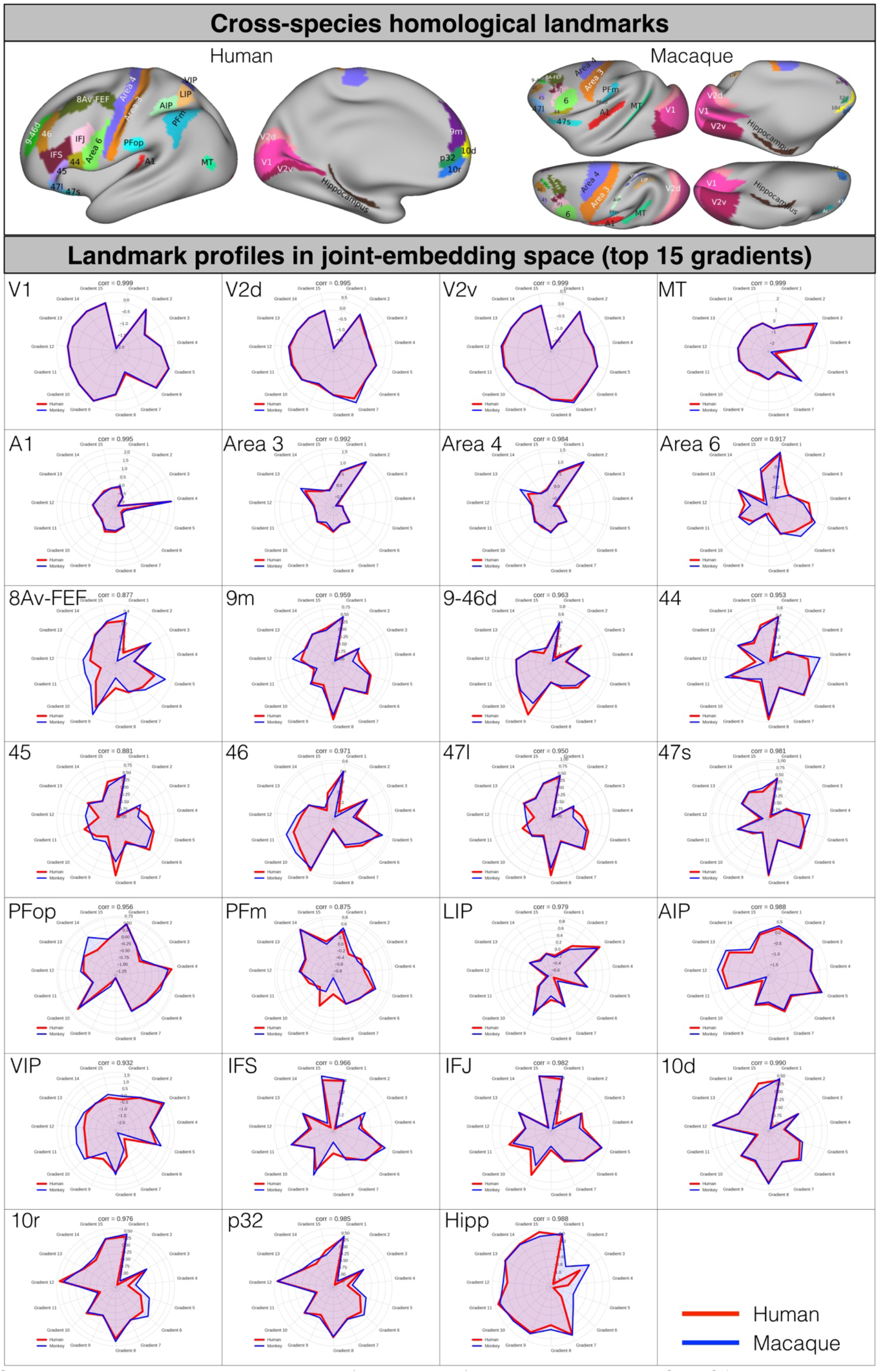
Cross-species homologous landmarks (upper panel) and the gradient profile of 27 landmark regions in radar plots for human and macaque monkey (lower panel). The top 15 joint-embedding gradients were used for profiling the cross-species common species.

**Figure S2.**
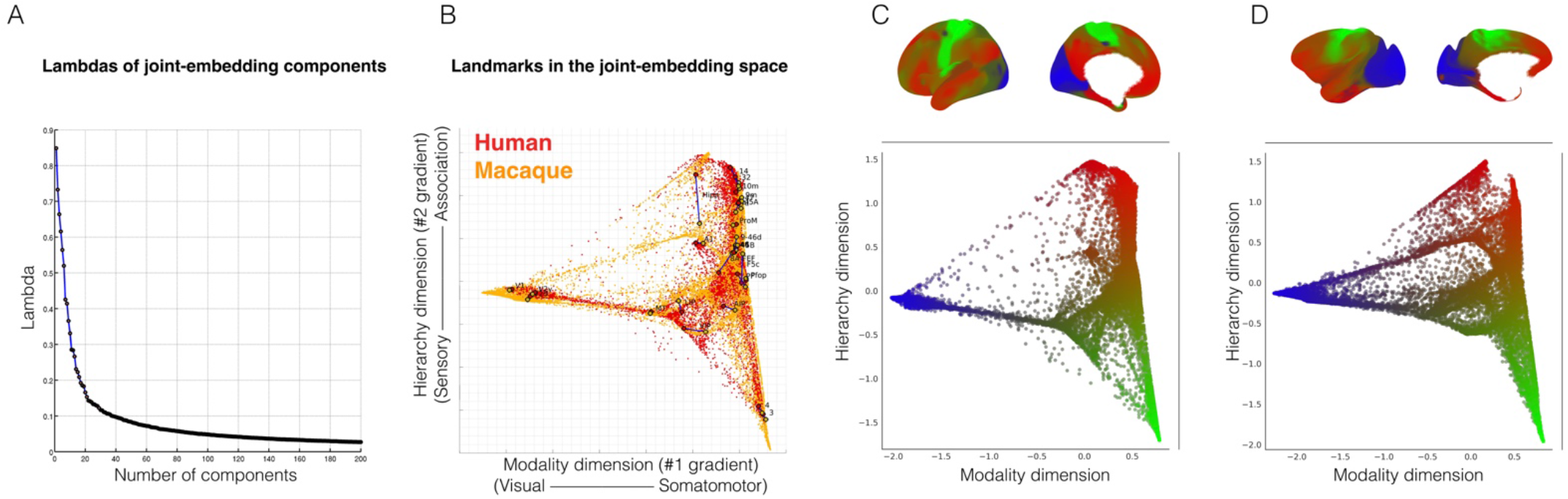
A) The lambdas (i.e. eigenvalues) of the joint-embedding at 1% sparsity threshold of the connectivity matrix within species. B) The first two gradients of the joint-embedding space represent the modality and the hierarchy dimensions. The homologous landmark pairs are close in the joint-embedding space. C) The gradients in joint-embedding space and presented on human cortical surface. D) The gradients in joint-embedding space and presented on macaque cortical surface. The gradients in joint-embedding space are color-coded in blue in visual cortex, green in somatomotor cortex and red at the apex of hierarchy dimension, the association cortex in both human and macaque monkey.

**Figure S3.**
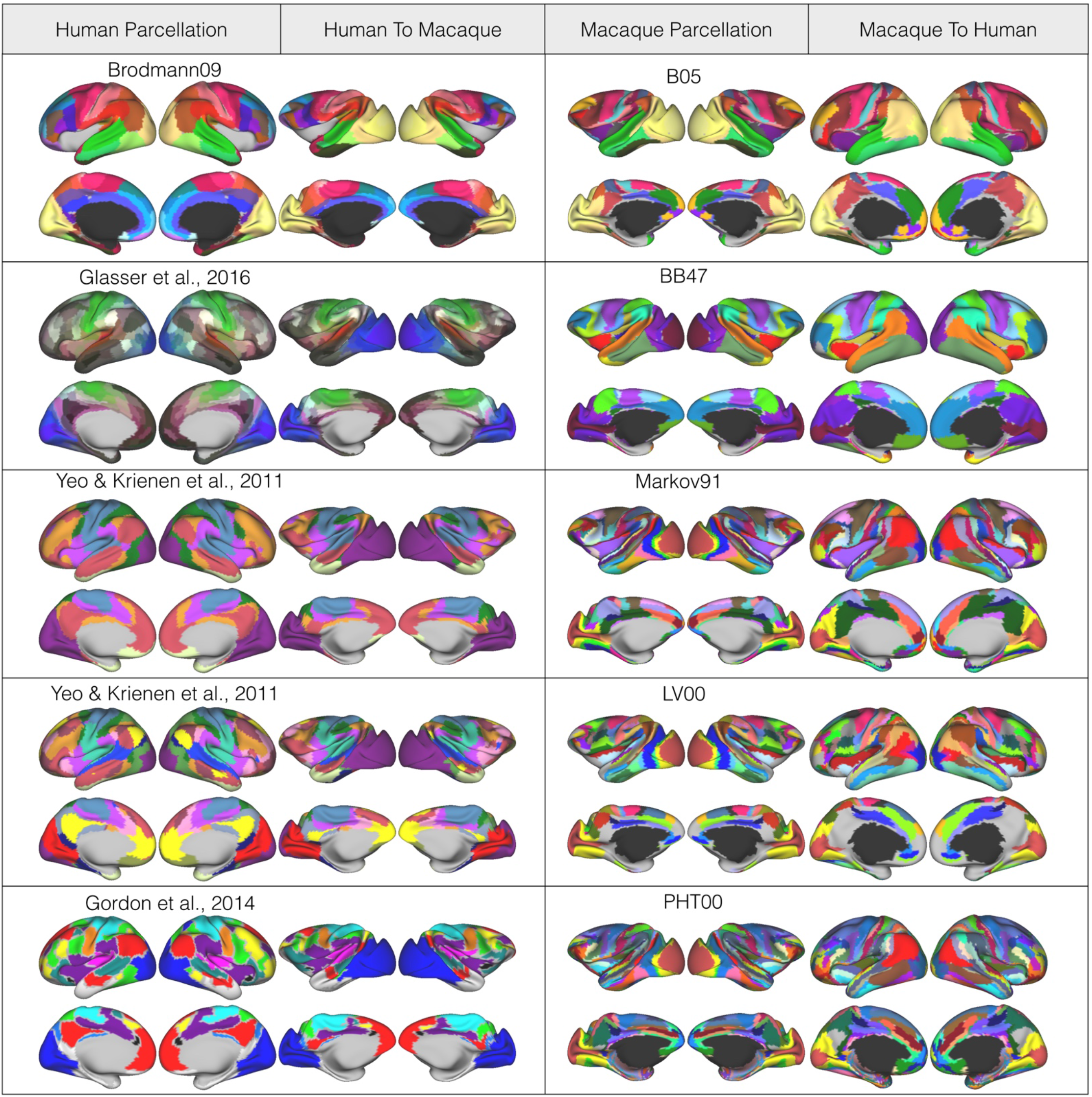
Mappings of various cortical parcellations based on the established cross-species cortical alignment in MSM. Left: The established human parcellations and the corresponding human-to-macaque parcellations. Right: The established macaque parcellations and the corresponding macaque-to-human parcellations.

**Figure S4.**
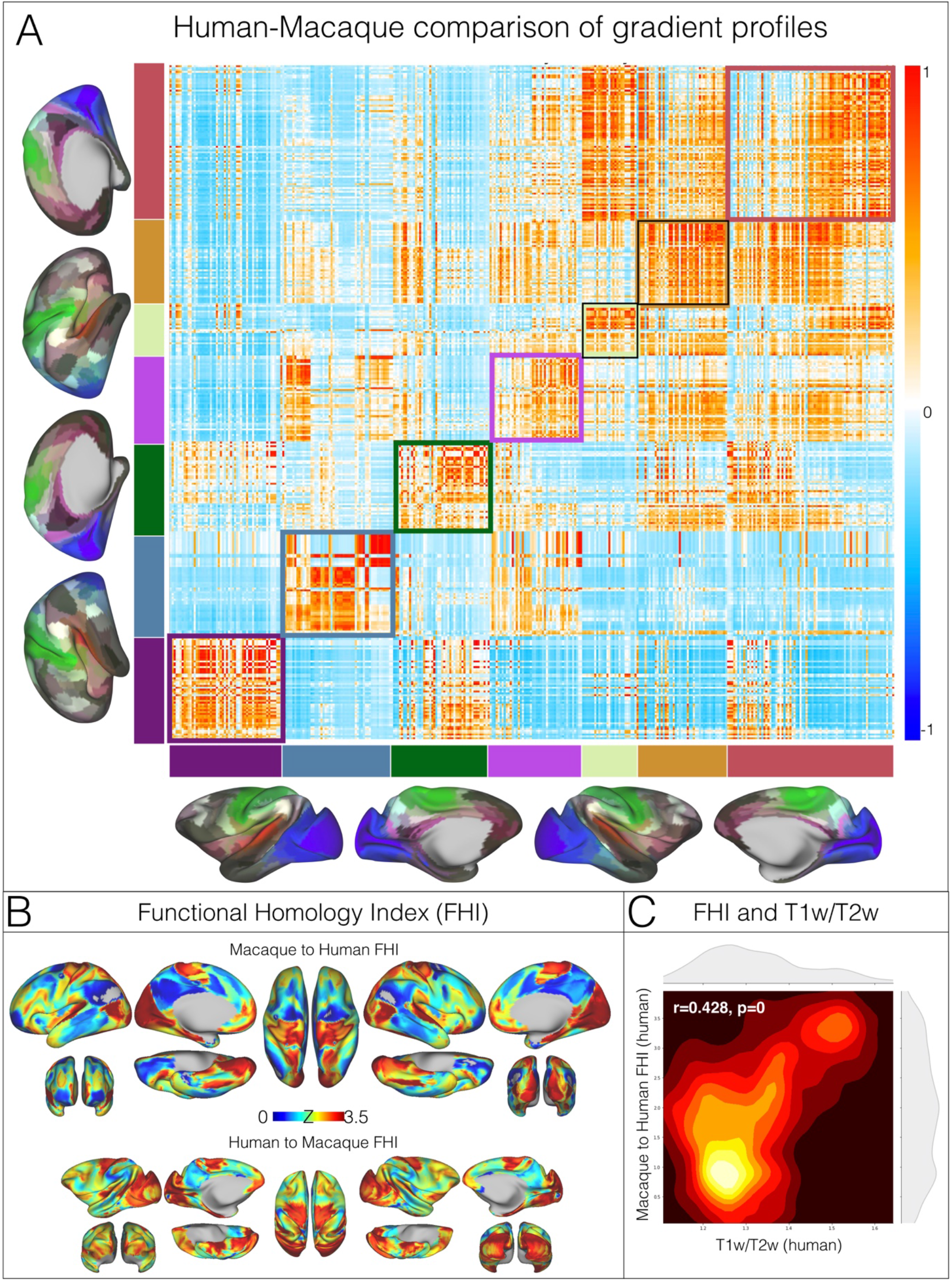
Pairwise cross-species comparison based on the functional gradient profiles and the functional homology index in human and macaque monkey. A) The parcel-wise cross-species similarity matrix parcellated by Glasser parcellation in human and its aligned human-to-macaque for macaque. The parcel-wise cosine similarity is averaged within the parcel pairs and ordered by Yeo2011 network identity. The within network shows greater similarity than between networks between macaque and human. B) The macaque-to-human (upper) and the human-to-macaque (bottom) functional homology maps calculated as the local maximal similarity of functional gradient profiles across species. The maps indicate the maximal likelihood that the functional organization in one species can be represented in other species. C) The spatial pattern of functional homology index is significantly associated with the T1w/T2w map in human.

**Figure S5.**
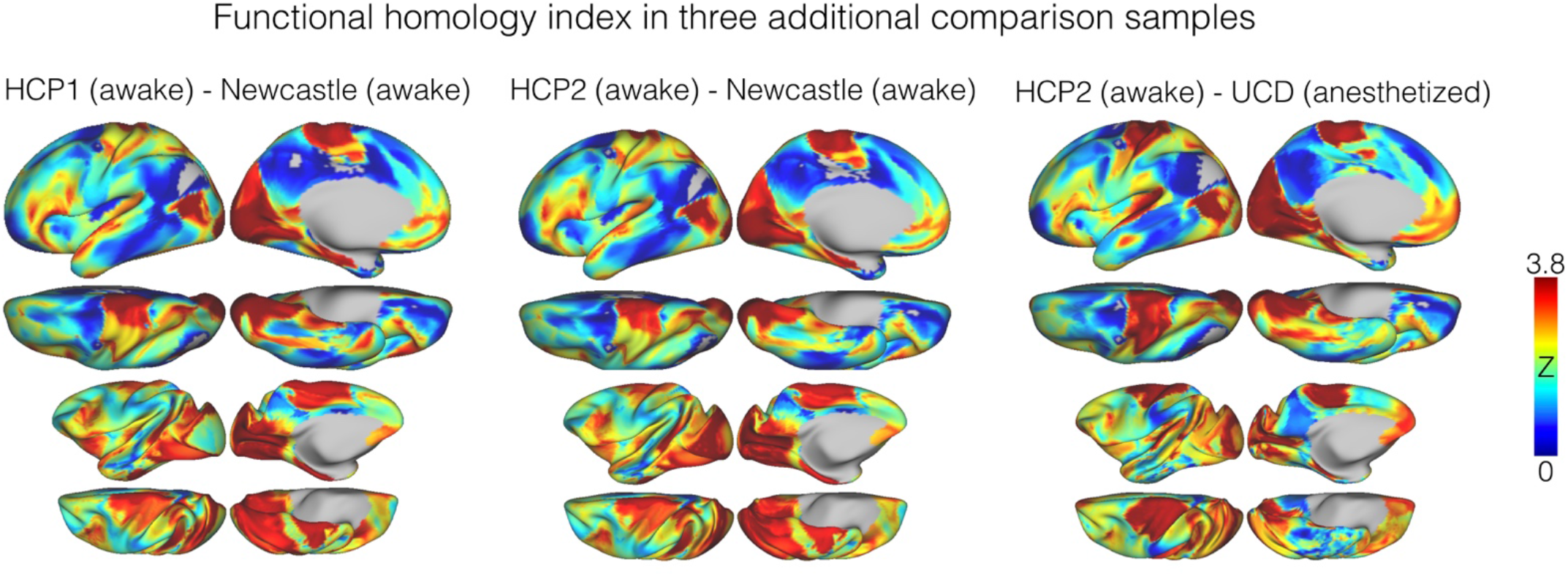
The functional homology map established based on the joint-embedding approach can be replicated in comparison samples.

**Figure S6.**
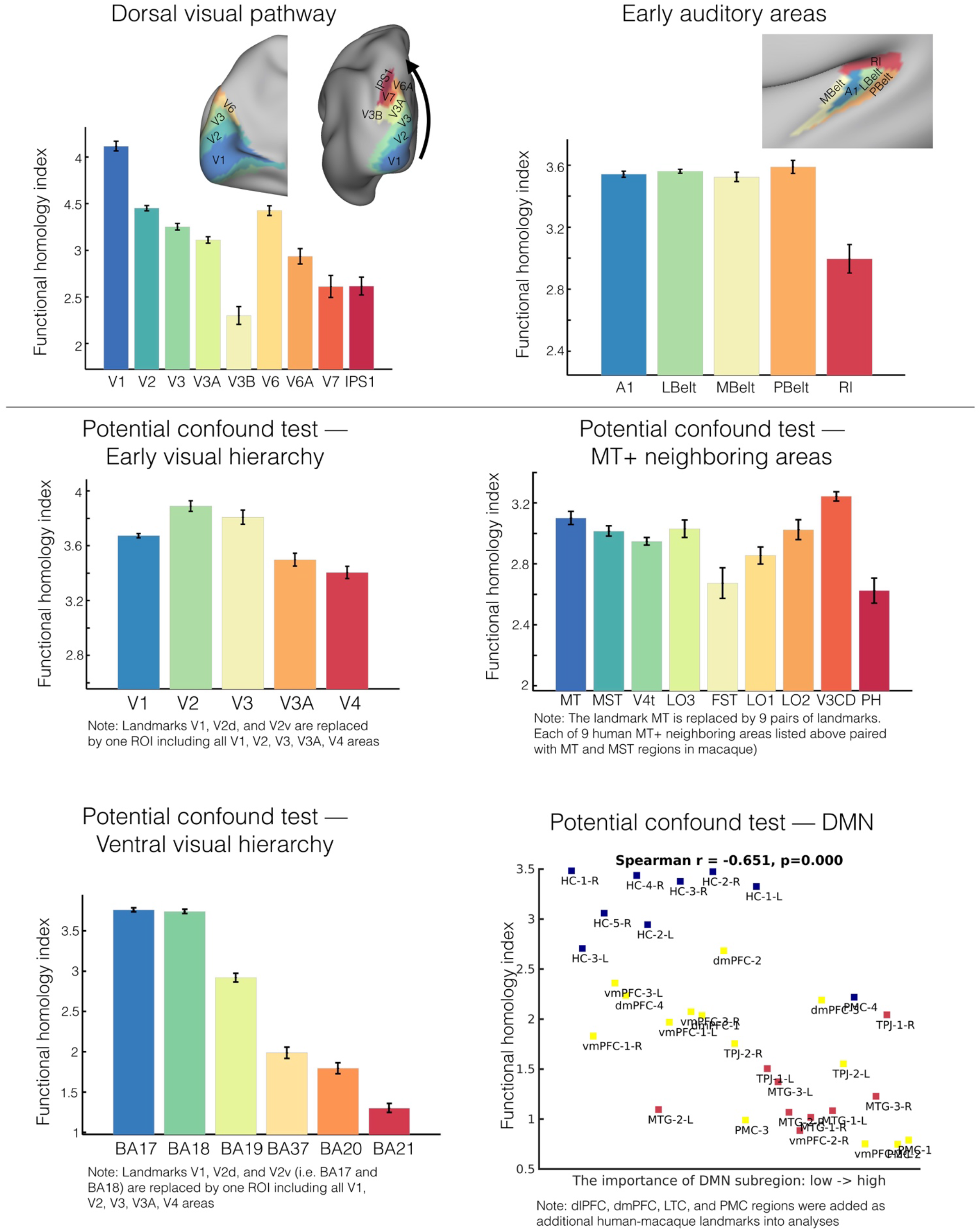
Functional homology in dorsal visual and primary auditory systems (upper panel). Potential confounds were tested to address whether the hierarchy patterns of function homology in local visual territories are driven by the specific landmarks in alignment. The general principle for testing was to include broader neighboring regions used in the hierarchy test and repeated the analyses. The hierarchical tests remained largely intact in the early visual, ventral visual systems, but MT+ neighboring areas. The association between FHI and the importance of DMN subregions was relatively unchanged when including dlPFC, dmPFC, LTC and PMC in the landmark set.

