## Supplemental Materials for "Cross-species Functional Alignment Reveals Evolutionary Hierarchy Within the Connectome"

### Methods

**Data.** Human and macaque monkey datasets in this study were from openly available sources. The human dataset was selected from the unrelated participants of the Human Connectome Project (HCP, https://www.humanconnectome.org)[1]. The macaque data stemmed from the recently established Nonhuman Primate sharing consortium PRIME-DE (http://fcon_1000.projects.nitrc.org/indi/indiPRIME.html)[2].

**Macaque monkey.** Three cohorts of macaque samples from PRIME-DE have been included in the present study. The data was preprocessed as described in our previous work[3,4].

**Oxford data (anesthetized).** The full dataset consisted of 20 rhesus macaque monkeys (macaca mulatta) scanned on a 3T with a 4-channel coil [5]. The resting-state fMRI (R-fMRI) data were collected while the animals were under anesthesia with 2 mm isotropic resolution, TR=2s, 53.3 min (1600 volumes). No contrast-agent was used during the scans. Nineteen macaques with successful preprocessing and surface reconstruction were included in the present study (all males, age=4.01+/-0.98, weight=6.61+/-2.04).

**UC-Davis data (anesthetized).** The full dataset consisted of 19 rhesus macaque monkeys (macaca mulatta) scanned on a Siemens Skyra 3T with a 4-channel clamshell coil. The resting-state fMRI data were collected with 1.4x1.4x1.4mm resolution, TR=1.6s, 6.67 min (250 volumes) under anesthesia. No contrast-agent was used during the scans. Nineteen macaques were included in the present study (all female, age=20.38+/-0.93, weight=9.70+/-1.58).

**Newcastle data (awake).** The full data set consisted of 14 rhesus macaque monkeys (macaca mulatta) scanned on a Vertical Bruker 4.7T primate dedicated scanner[6–12]. We included 10 animals (8 males, age=8.28+/-2.33, weight=11.76+/-3.38) who were scanned awake. The fMRI session was acquired with 1.2x1.2x1.2mm resolution, TR=2s, 8.33-min per scan (250 volumes x 2 scan) for each animal. No contrast-agent was used during the scans.

**Human.** We selected the R-fMRI data from the unrelated participants in the HCP S500 release[1]. The first R-fMRI scan acquired on day one has been included in the current analysis, containing a 15-min run (phase encoding left-right) for each participant. The details of the acquisition and the preprocessing can be found at https://www.humanconnectome.org/study/hcp-young-adult/data-releases. We randomly split the human data into two subsets (subset HCP1, n=93, 46 females, age=29.23+/-3.49; subset HCP2, n=94, 36 females, age=28.99+/- 3.43). These two subsets were grouped into two human and anesthetized macaque comparisons (HCP1-Oxford, HCP2-UCD) and two human and awake macaque comparisons (HCP1-Newcastle, HCP2-Newcastle).

**Preprocessing.** The macaque monkey data were preprocessed using the customized HCP-like pipeline from DAF's laboratory and the Computational Connectome System[13]. The details of the data preprocessing were described previously[3,4]. Briefly, the R-fMRI data were preprocessed including temporal compression, motion correction, 4D global scaling, nuisance regression using white matter (WM), and cerebrospinal fluid (CSF) signal and Friston-24 parameter models, bandpass filtering (0.01-0.1Hz), detrending and co-registration to the native anatomical space. The data were then projected to the native mid-cortical surface and smoothed along the surface with FHWM=3mm. Finally, the preprocessed data were down-sampled to a 10k (10,242 vertices) resolution surface. Similar with the macaque preprocessing, the human data have been minimally preprocessed in the HCP pipeline in addition with the bandpass filtering (0.01-0.1Hz), spatial smoothing along the surface (FWHM=6mm) and downsampling to the 10k (10,242 vertices) mid-cortical surface[14,15].

**Cross-species landmarks.** The landmarks were selected based on the milestone study from Van Essen' group[16] and recent cross-species comparison based fMRI from Mars' group[17–19]. Only potential landmarks that have been reported in at least two studies were included in the current work. The final set included 27 landmarks (Table S1). The area definition in human was based upon the most recent multi-modal human parcellation[20]. For the landmark area in macaque, we first collected the area definitions from seven macaque atlases and used the vertices that at least overlapped within two atlases for the final macaque landmarks[21–28]. The details of the studies used to define the landmarks and the atlas references were listed in Table S1.

**Joint-embedding**

In previous work on manifold alignment, spectral embedding (e.g. diffusion maps) has demonstrated the ability to align the connectivity structure across individuals[29–32]. Recently, this approach has been used to characterize the connectivity topographies and capture the cortical gradients spanning along the unimodal (visual and somatomotor cortices) and transmodal regions (association cortex) within each species in human and macaque monkey[33,34]. Here, in order to align human and macaque monkey cortex, the challenge is to extract comparable cross-species components, rather than applying embeddings for each species individual and subsequently performing component matching. To address this challenge, we propose a joint-embedding approach to compute matched components (referred to as 'gradients') in human and macaque monkey.

First, we constructed a joint similarity matrix by concatenating within- and cross-species similarities of connectivity patterns (Fig 1A), as defined in

$W = [W_{human}, W_{human\ to\ monkey}\ ;\ W_{macaque\ to\ human}, W_{monkey}]$.

The diagonal within-species similarity matrices ($W_{human}$ and $W_{monkey}$) are calculated using cosine similarity of row thresholded functional connectivity at each vertex in each species[33]. The functional connectivity was calculated at the group-level by averaging the individual connectivity matrix first within each of the comparison samples. The off-diagonal cross-species similarity matrix $W_{human\_to\_monkey}$ (and its transpose $W_{monkey\_to\_human}$) was calculated based on the landmark similarity profile of the functional connectivity pattern. Specifically, similar to a previous study from

Mars [35], we first computed the thresholded vertex-to-vertex connectivity matrix ($C_{human}$ and $C_{monkey}$) and averaged the vertex-wise connectivity to each landmark respectively to generate the vertex-to-landmark connectivity matrix ($L_{human}$ and $L_{monkey}$) for each species. Based on these two connectivity matrices profile, we calculated the vertex-to-landmarks similarity matrix ($S_{human}$ and $S_{monkey}$) within each species. That is, for each vertex within a species, the row *i* of matrix *S* is defined as the cosine similarity between row i of C and row i of L. Note that the 27 landmarks were matched homologous areas between human and macaque monkey, in other words, the columns of $S_{human}$ and $S_{monkey}$ are matched. Then we measured the cross-species similarity matrix $W_{human\_to\_monkey}$ (and its transpose $W_{monkey\_to\_human}$) by comparing the similarity pattern to the homologous landmarks across species. To determine the threshold for the connectivity matrix within each species, we tested the sparsity thresholds at 1% to 10% and examined the distance of matched homologous landmarks between human and macaque in the resultant gradient space. The sparsity threshold 1% generated the most similar cross-species gradients and was employed in the final analysis.

Next, we applied the diffusion embedding algorithm on the concatenated matrix W, resulting in a set of components[31]. Of note, the joint similarity matrix W is a symmetric matrix across species. The diagonal block matrices contain the within species connectivity profiles in human and macaque, encoding backbone connectivity structure (thresholded at top 1%), while the off-diagonal matrices provided a coupling across species via the comparable landmarks. Therefore, for each of the obtained components, the first half of entries correspond the human vertices and the second half macaque vertices (Fig 1A). Each component provides a set of matched cortical gradients covering the human and macaque cortices, which can be served as one of the dimensions of the common cross-species coordinate space. We first extracted the top 200 components and selected only the top k components to construct a gradient pool for the following surface matching procedure. Here, k is determined as the inflection point of eigenvalues (lambdas) on the scree plot (Fig S2A). Twenty-five components (i.e. gradients) were selected in the HCP1-Oxford comparison sample (21 for HCP1-Newcastle, 18 for HCP2-UCDavis, 21 for HCP2-Newcastle).

Finally, we used the gradients from the above gradient pool as the surface features and aligned the human and macaque cortical surface with Multimodal Surface Matching (MSM)[36]. In order to avoid misalignment in the medial wall between human and macaque, we added the medial wall mask as an additional feature into MSM. The MSM configuration parameters 'config_MSMsulc_pairwise' was used in alignment. To optimize the number of gradients for the final alignment, we entered the top 5, 10, 15, and 20 gradients in MSM and determined the performance using 27 landmarks labels as the inspection standard. Top 15 components were selected for the final alignment in all four comparison samples. Accordingly, these 15 components were used as gradient profiles to build the common coordinate space between human and macaque monkey. We examined the alignment performance by applying the surface deformation to the myelin sensitive maps (i.e. T1w/T2w) and compared the aligned myelin prediction map with the actual T1w/T2w estimation in aligned species (Fig S4C). In addition, several well-established human and macaque parcellations and networks can be registered well from human to macaque, vice versa (Fig S3). We also calculated the cross-species similarity matrix based on 15 gradients profiles at each vertex and demonstrated the parcel-wise similarity matrix using the most recent multimodal parcellations for human and its aligned human-to-macaque for macaque (Fig S4). It can be seen that in general the cross-species similarity revealed that greater similarity within network than between networks (Fig S4A).

**Functional Homology Index (FHI)**

In order to quantify cross-species regional similarities of functional organization in the functional common space, we further developed the *Functional Homology Index* (FHI, Fig 2A). Specifically, for each pair of coordinates identified as corresponding between species in MSM, we quantified the maximum cosine similarity of 15 gradients as FHI across species within corresponding searchlights (radius = 12 mm on the midthickness surface). The searchlight approach mitigates the possibility of excessive topological constraints from MSM, while limiting the identification of matches that are unfeasible. The maximum similarity within the corresponding searchlight quantified the highest likelihood that the functional gradients at each vertex in human can be represented in macaque (Fig 2B) and vice versa (Fig S4B).

**The activation possibility strength of BrainMap cognitive component**

To quantify the relationship between functional homology and cognitive function, we employed a similar analysis as described in recent studies [33,37]. The human cognitive functions were represented using the activation possibility maps of 12 cognitive components from a previous large-scale meta-analysis based on the BrainMap database [38]. Specifically, we first grouped the macaque-to-human FHI map into 10-percentile bins. For each of 12 cognitive components, the activation strength was normalized by dividing the sum of each components' activation possibility and then sum within each of the 10 bins. The score in the heatmap represents the total activation possibility associated with a given cognitive component within each of the 10-percentile bin regions. The cognitive components were ordered based on the activation strength weighted by the log scale of percentile.

**Evolutionary Deformation and Area Expansion**

The evolutionary surface expansion was calculated at each vertex based on the correspondence established in MSM. Specifically, we first estimated the vertex-wise surface area of the 32k standard surface mesh in native space for each of human and macaque individuals. We then resampled and smoothed (FWHM=6mm in human and FWHM=3mm in macaque) the area estimations to 10k surface using areal interpolation [39]. Next, the individual area maps were averaged across all the individual to generate area map for each of human (n=187) and macaque samples (n=48). After that, we estimated the macaque surface area at each of corresponding human vertices using the registration sphere in MSM [39]. The final relative area expansion was calculated by dividing the human surface area by the macaque surface area at each vertex on human surface. Similarly, we calculated the relative area at each vertex on macaque surface, suggesting the starting points of the expansion origin from macaque to human.

To further demonstrate the evolutionary direction on the surface, we calculated macaque-to-human deformation vectors. To facilitate the visualization in highly folded regions (e.g. insular), we used the 'very_inflated' surface for both human and macaque monkey. Specifically, we identified the macaque-to-human coordinates for each of the vertices corresponding to the very_inflated macaque surface using MSM registration sphere. Next, we calculated the vector based on the MSM aligned coordinates from macaque to human. The length of the vector represents the strength of the evolutionary deformation along the direction from macaque to human surface.

**Table S1**. The landmarks definition and the atlas used that generated the landmarks

| # | Cross-species comparison studies | Area name in Human | Label name in human atlas (Glasser et al., 2016) | Area name in Macaque | Label name in macaque atlases | Macaque Atlas Reference |
|---|---|---|---|---|---|---|
| 1 | Neubert et al., 2014, Glasser et al., 2016 | Ventro-Medial Prefrontal Cortex | 10r | VMPFC/Area14 | 14 | Markov et al., 2011; 2014 |
| 2 | Neubert et al., 2014; Glasser et al., 2016 | Area 32 | p32 | Area 32 | 32 | Markov et al., 2011; 2014; Ferry et al., 2000 |
| 3 | Neubert et al., 2014; Sallet et al., 2013; Glasser et al., 2016 | Area 9 medial | 9m | Area 9 medial | 9m | Preuss and Goldman-Rakic 1991; Paxinos and Franklin 2000 |
| 4 | Neubert et al., 2014; Sallet al., 2013; Glasser et al., 2016 | Area 9/46 dorsal | 9-46d | Area 9/46 dorsal | 9/46d | Markov 2011; 2014; Paxinos and Franklin 2000 |
| 5 | Neubert et al., 2014; Sallet al., 2013; Petrides, 2015; Glasser et al., 2016, Van Essen 2007; Van Essen and Dierker 2007 | Area 8A/Frontal Eye Field | 8Av, FEF | Area 8A, FEF | 8A, FEF | Van Essen 2007; Lewis and Van Essen 2000; |
| 6 | Neubert et al., 2014; Mars et al., 2011; Glasser et al., 2016 | Area PFop | PFop | Area PFop | PFop | Paxinos and Franklin 2000 |
| 7 | Neubert et al., 2014; Mars et al., 2011; Glasser et al., 2016 | Anterial Intraparietal | AIP | Anterior Intraparietal Area | AIP | Markov et al., 2011; 2014 |

| | | | | | | |
|---|---|---|---|---|---|---|
| | | Area | | | | |
| 8 | Neubert et al., 2014; Mars et al., 2011; Glasser et al., 2016 | Ventral Intraparietal Area | VIP | Ventral Intraparietal Area | VIP | Felleman and Van Essen 1991; Markov et al., 2011; 2014 |
| 9 | Neubert et al., 2014; Mars et al., 2011; Glasser et al., 2016 | Mid-Intraparietal Sulcus | LIPd, LIPv | Mid-Intraparietal Sulcus | LIP | Felleman and Van Essen 1991; Markov et al., 2011; 2014 |
| 10 | Neubert et al., 2014; Mars et al., 2011; Glasser et al., 2016 | Mid-Inferior Parietal Lobule | PFm | Area PF | PF | Paxinos and Franklin 2000 |
| 11 | Neubert et al., 2014; Glasser et al., 2016 | Area 6 ventral, Area 6 dorsal | 6v, 6r | Area F5c, Area F5a | F5c | Markov et al., 2011; 2014 |
| 12 | Neubert et al., 2014; Glasser et al., 2016 | Inferiro Frontal Sulcus | IFSp, IFSa | Area 45B | 45B | Paxinos and Franklin 2000 |
| 13 | Neubert et al., 2014; Glasser et al., 2016 | Inferior Frontal Junction | IFJa, IFJp | Area 44 | 44 | Markov et al., 2011; 2014 |
| 14 | Neubert et al., 2014; Glasser et al., 2016 | Area 44 ventral | 44 | Area ProM | ProM | Markov et al., 2011; 2014 |
| 15 | Neubert et al., 2014; Glasser et al., 2016 | Area 45 | 45 | Area 45A | 45A | Markov 2011; 2014; Paxinos and Franklin 2000 |
| 16 | Neubert et al., 2014; Glasser et al., 2016 | Area 47 | 47l | Area 47 | 47 | Paxinos and Franklin 2000 |

| | | | | | | |
|---|---|---|---|---|---|---|
| 17 | Neubert et al., 2014; Glasser et al., 2016 | Operculum Frontale | 47s | Anterior Insula | AI | Paxinos and Franklin 2000 |
| 18 | Neubert et al., 2014; Glasser et al., 2016 | Area 46 | 46 | Area 46 | 46 | Felleman and Van Essen 1991 |
| 19 | Neubert et al., 2014; Glasser et al., 2016 | Mid-Inferior Parietal Lubule | 10d | Area 10m | 10m | Ferry et al., 2000 |
| 20 | Van Essen 2007; Van Essen and Dierker 2007; Glasser et al., 2016 | Area V1 | V1 | Area V1 | V1 | Van Essen 2007; Markov 2011; 2014 |
| 21 | Van Essen 2007; Van Essen and Dierker 2007; Glasser et al., 2016 | Area V2 dorsal | V2d | Area V2 dorsal | V2d | Van Essen 2007; Markov 2011; 2014 |
| 22 | Van Essen 2007; Van Essen and Dierker 2007; Glasser et al., 2016 | Area V2 ventral | V2v | Area V2 ventral | V2v | Van Essen 2007; Markov 2011; 2014 |
| 23 | Van Essen 2007; Van Essen and Dierker 2007; Glasser et al., 2016 | Area A1 (Auditory) | A1 | Area A1 (Auditory) | A1 | Van Essen 2007; Markov 2011; 2014 |
| 24 | Van Essen 2007; Van Essen and Dierker 2007; Glasser et al., 2016 | Area MT | MT | Area MT | MT | Van Essen 2007; Markov 2011; 2014 |
| 25 | Van Essen 2007; Van Essen and Dierker 2007; Glasser et al., 2016 | Area 3 | 3a, 3b | Area 3 | 3 | Van Essen 2007; Markov 2011; 2014 |

| | | | | | | |
|---|---|---|---|---|---|---|
| 26 | Van Essen 2007; Van Essen and Dierker 2007; Glasser et al., 2016 | Area 4 | 4 | Area 4 | 4 | Van Essen 2007; Markov 2011; 2014 |
| 27 | Van Essen 2007; Van Essen and Dierker 2007; Glasser et al., 2016 | Area Hippocampus | H | Area Hippocampus | Hipp | Van Essen 2007; Markov 2011; 2014 |
